# Accurate brain age prediction using recurrent slice-based networks

**DOI:** 10.1101/2020.08.04.235069

**Authors:** Pradeep K. Lam, Vigneshwaran Santhalingam, Parth Suresh, Rahul Baboota, Alyssa H. Zhu, Sophia I. Thomopoulos, Neda Jahanshad, Paul M. Thompson

## Abstract

BrainAge (a subject’s apparent age predicted from neuroimaging data) is an important biomarker of brain aging. The deviation of BrainAge from true age has been associated with psychiatric and neurological disease, and has proven effective in predicting conversion from mild cognitive impairment (MCI) to dementia. Conventionally, 3D convolutional neural networks and their variants are used for brain age prediction. However, these networks have a larger number of parameters and take longer to train than their 2D counterparts. Here we propose a 2D slice-based recurrent neural network model, which takes in an ordered sequence of sagittal slices as input to predict the brain age. The model consists of two components: a 2D convolutional neural network (CNN), which encodes the relevant features from the slices, and a recurrent neural network (RNN) that learns the relationship between slices. We compare our method to other recently proposed methods, including 3D deep convolutional regression networks, information theoretic models, and bag-of-features (BoF) models (such as BagNet) - where the classification is based on the occurrences of local features, without taking into consideration their global spatial ordering. In our experiments, our proposed model performs comparably to, or better than, the current state of the art models, with nearly half the number of parameters and a lower convergence time.

## 1. INTRODUCTION

In recent years, deep learning methods have gained considerable attention from the neuroimaging community, in many cases outperforming traditional methods for tasks such as segmentation, disease classification, and quality control^1–3^.

However, 3D neuroimaging data poses several challenges for the development of robust models. These models often require a large number of training parameters^4^ and the lack of training data relative to the number of parameters can make networks prone to overfitting^5^. In addition, the lack of models that have been pre-trained with large-scale 3D imaging datasets inhibits effective transfer learning. Networks pretrained with ImageNet have been shown to learn more generalizable features^6^ and networks trained with 2D slices have shown significant performance improvements on BrainAge estimation and Alzheimer’s disease classification tasks^7–8^. Logistically, the number of parameters in these models requires large GPU memory and the models typically take a long time to train.

In this paper, we propose a slice-based recurrent neural network (RNN) model, which uses a 2D convolutional neural network to encode information from sagittal slices of 3D brain images and a recurrent neural network to learn the inter-slice relationship. We apply this model to the task of BrainAge prediction and assess the models’ performance relative to several popular 3D CNN BrainAge prediction methods.

Briefly, the task of predicting a subject’s age from their brain scan has been attempted using a broad variety of methods^9^. Briefly, a large dataset of brain MRI scans - usually from healthy individuals free from neurological or psychiatric disease - is used as input to a machine learning algorithm that is trained to predict each person’s chronological age from their image, or from a set of features that have been derived from it (such as measures of cortical gray matter thickness in regions of interest). The methods used can be categorized into (1) classical machine learning methods, such as ridge regression and support vector machines^10^, and (2) deep learning methods, such as the CNNs evaluated here, that distil successively more abstract features from the raw images. There are also more complex methods that combine information from several types of brain imaging modalities (anatomical, diffusion-weighted, and functional MRI^11^). Here, we focused on the intermediate case, using a single data modality (anatomical MRI), which is the most commonly collected type of MRI and avoids the need to collect multiple MRI data modalities, some of which may not be generally available.

In our evaluations below, our model outperformed several popular models on an independent test set and outperforms several recently proposed state of the art methods, with a mean absolute error (MAE) of 2.8 years, and with only 1 million parameters. The model code and weights are freely available from our GitHub repository: https://github.com/USC-IGC/RNN_Slice_BrainAge

## 2. METHODS

### 2.1 Description of participants

We analyzed a brain MRI dataset from 16,356 individuals from the UK Biobank^12^. The UK Biobank is a large epidemiological study of 500,000 people residing in the UK, some of whom received neuroimaging. From the initial dataset of those with neuroimaging, we selected a subset of 10,446 who had no indication of neurological pathology, and no psychiatric diagnosis as defined by ICD-10 criteria. The age range of our dataset was 45-81 years. The entire dataset has a mean age of 62.64 (SD, 7.41; 47% women/53% men).

### 2.2 Preprocessing of scans

All scans were evaluated with a manual quality control procedure, where scans with severe artifacts were discarded. The remaining scans were processed using a standard preprocessing pipeline with nonparametric intensity normalization for bias field correction^13^ and brain extraction using FreeSurfer^14^ and linear registration to a (2 mm)^3^ UKBB minimum deformation template using FSL FLIRT^15^. The final dimension of the registered images was 91 × 109 × 91.

### 2.3 Dividing into test

Data was randomly divided into 3 non-overlapping sets: (1) 7,312 scans were used for training (mean age: 62.68; 7.41 std); (2) 940 scans were used as the validation set (mean age: 62.59; 7.48 std) to fine-tune hyperparameters and to determine the model stopping point; and (3) 2,194 scans were set aside for independent testing of the trained model (mean age: 62.50, 7.41 std).

### 2.4 2D slice sequence LSTM

Our model (**Figure 1**) contains two main components: a 2D convolutional network that encodes relevant intra-slice features and a recurrent network that processes an ordered series of slice encodings and outputs a single age embedding for the scan (Consisting of 91 slices for the sagittal plane of section). This embedding is fed into a simple one-layer dense network to predict age.

**Figure 1.**
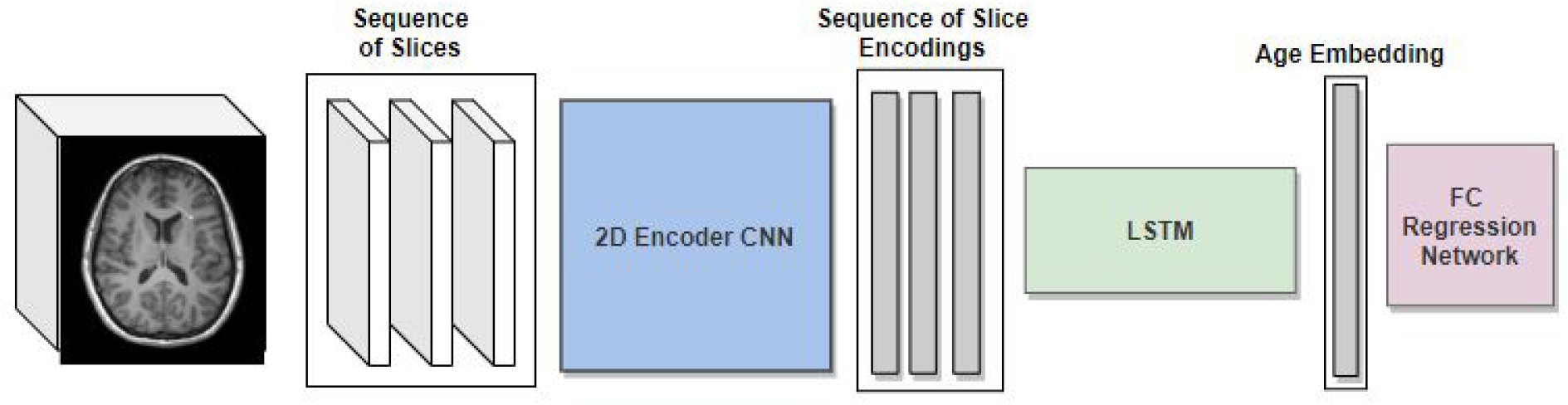
The architecture of our model is shown here. Each 3D brain MRI scan (*left*) is converted to a sequence of 2D slices before feeding these to the 2D CNN Encoder, which produces the sequence of encodings for the slices. The LSTM takes this sequence as input, and learns the relationship between the slices and produces an output embedding, which is fed into a Fully Connected Regression Network (*right panel*). This predicts the Brain Age.

Our convolutional encoder network follows a very similar architecture to Peng et al.^4^ The 2 major differences are that we replace BatchNorm with InstanceNorm and replace all 3D convolutions/pooling operations with their 2D counterparts. The network (**Figure 2**) has seven blocks. The first five blocks consist of a 3×3 2D convolutional layer (stride=1, padding=1) followed by an instance norm^16^ layer, a 2×2 max-pool layer (stride=2), and a ReLU activation. The number of filters in the first block is 32 - and doubles until 256 - with both layers 4 and 5 having 256 filters. The sixth block contains a 1×1 2D convolutional layer (stride=1), followed by an instance norm layer, and a ReLU activation. The convolution layer contains 64 filters. The seventh block contains an average pooling layer, a dropout layer (set to p=0.5, during training time), and a 1×1 2D convolutional layer with stride 1. The number of filters in the final convolutional layer is determined by the dimensionality of the per slice feature encodings (n=2 for our experiments).

**Figure 2.**
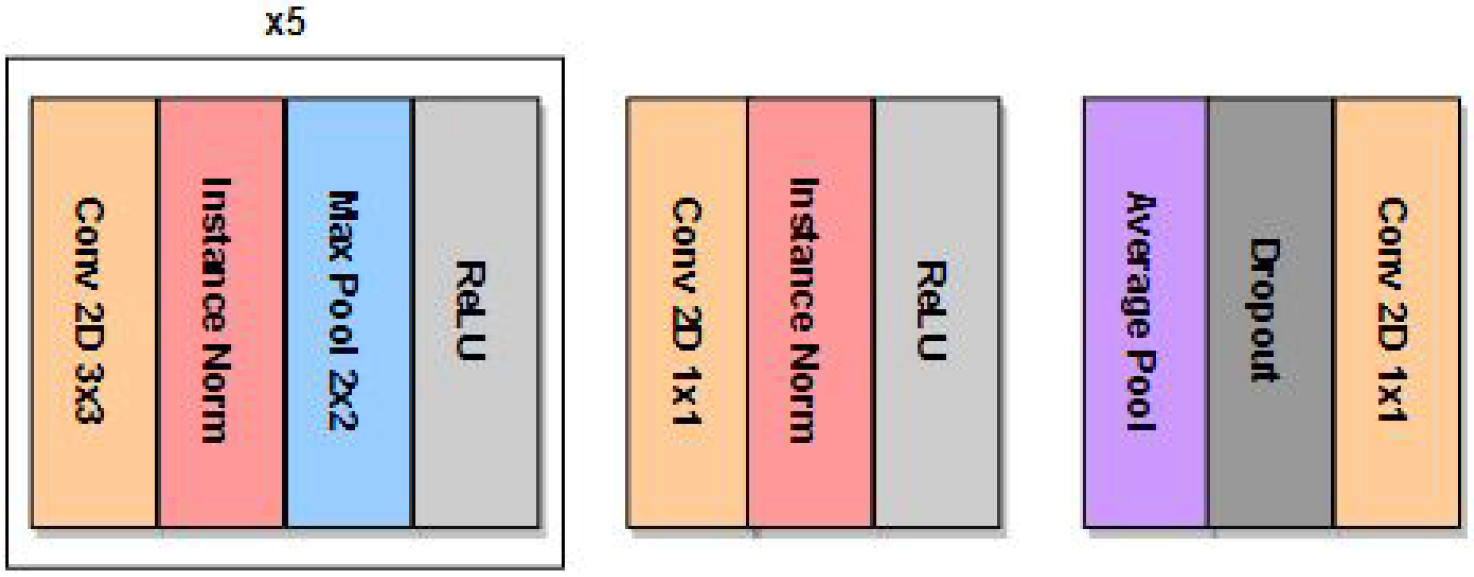
The architecture of our core encoder network is presented here. The model takes 2D brain MRI slices and contains 7 blocks. Each of the first 5 consecutive blocks consists of a 3×3 2D convolution layer, a Instance Norm layer, a Max Pooling layer and a ReLU activation. The 6th block contains one 1×1 2D convolution layer, an Instance Norm layer and a ReLU activation. The 7th block contains an average pooling layer, a dropout layer, and a 1×1 2D convolution layer.

We use a recurrent network to learn a complete representation from slice-based encodings, treating all slices in a single scan as an ordered sequence. We employ a single layer Long-Short-Term-Memory (LSTM)^17^ network with a hidden state/cell state dimensionality of 128 with an input dimensionality of 2 (corresponding to the size of the per slice feature embeddings from the cnn encoder). The final hidden state of the LSTM is connected to a 64-node dense network with ReLU activation, which is then fed into a 1-node output layer.

We also compared our model against several popular deep learning models found in the literature, detailed below.

### 2.5 3D Regression CNN

Deep 3D convolutional regression networks have been used in several instances for brain age prediction^18–19^. These networks extend the traditional VGG and ResNet architectures to 3D images by replacing 2D convolution/maxpool operations with their 3D counterparts. For our comparison, we used the 3D variant of the 2D encoder CNN used in our sliced-based LSTM model; only replacing the 2-node feature embedding with a single node regression output.

### 2.6 3D “Info” CNN

Peng et al.^4^ framed brain age prediction as a soft-class classification problem. Here a 3D convolutional network is trained by fitting data to a discretized normal distribution with mean equal to true chronological age of the individual and a standard deviation of 1.

The network is trained by minimizing the KL Divergence (**Figure 3**) between the discrete true distribution and softmax network output with the same number of bins.

**Figure 3.**
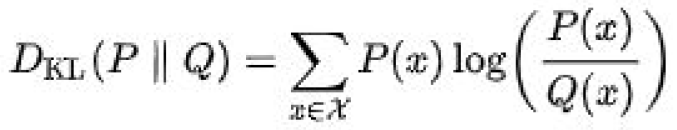
The equation for Kullback-Leibler (KL) Divergence is presented here, where P(x) is the discrete true distribution and Q(x) is the softmax network output.

Our implementation follows that of our regressor, except that it replaces the single node output with 37 nodes and adds a softmax layer to the end of the network.

### 2.7 3D BagNet

Pawlowski and Glocker^20^ adapt the BagNet model^21^ for brain age regression and sex classification from structural MRI.

BagNet is based on the bag-of-features (BoF) models where the classification is based on the occurrences of small local features without taking into consideration the global spatial ordering. This is done by limiting the receptive field of a ResNet50 model from 177 pixels to 9, 17, or 33 pixels. The smaller receptive field enforces locality in the extracted features.

For our comparison, we replicated the Pawlowski and Glocker^20^ Bagnet17 model (each imaging feature used in regression is based on a receptive field of 17) for age regression. This implementation uses the ResNet50 architecture but replaces 2D convolutions with 3D convolutions; it also replaces batch normalization with instance normalization, and cuts the number of feature maps in 2.

### 2.8 Training and optimization

Weights for all models are initialized using Kaiming weight initialization scheme^22^. All regression models were initialized with the bias of the final layer equal to the mean (62.68) of the training dataset.

All networks were trained using the Adam Optimizer^23^ with a batch size of 8, except for BagNet, which was run with batch size of 4 due to memory constraints and an initial learning rate of 1e-4. This learning rate is decayed by a factor of 0.8 based on the plateau of loss on the validation set. In addition, all models employed a gradient clipping of 1. All models optimized an MSE loss with the exception of the Info CNN which optimizes the KL divergence loss described above.

Training was capped at 80 epochs with early stop enabled for plateau of validation loss. Checkpoints used for evaluation were selected based on minimizing average loss on the validation dataset.

## 3. RESULTS

Models were evaluated on a non-overlapping, independent set of 2,194 scans from healthy individuals who had no indication of having brain pathology, and no medical diagnosis of psychiatric or neurological illness. Performance for different models may be found in **Table 1**.

**Table 1.** Each row represents the results obtained by the respective models. The last column,”Num param” represents the total number of parameters required by the model. It can be clearly seen from the results that our 2D slice sequence LSTM has the best performance compared to the other models, in addition to having the least number of parameters required to train the model.

| Method | RMSE | MAE | Corr | Resid Corr | Num param |
| --- | --- | --- | --- | --- | --- |
| 3D Reg CNN | 3.80 | 3.05 | 0.87 | -0.61 | 2,016,065 |
| 3D Info CNN | 3.83 | 3.11 | 0.87 | -0.69 | 2,018,000 |
| 3D BagNet17 | 3.79 | 3.00 | 0.85 | -0.53 | 15,734,977 |
| 2D Slice | **3.61** | **2.86** | 0.87 | **-0.33** | **1,068,065** |

*Corr* denotes the correlation between true age and predicted age in the unseen test data: higher values are better, denoting more accurate predictions. *Resid Corr* denotes the residual correlation between true age and the difference between a subject’s predicted age and true age - an undesirable effect of regression toward the mean, that has been noted in prior works by Smith^11^, Cole et al.^16^, and others: lower values are better.

The recurrent slice-based method obtains the lowest RMSE, MAE, and Residual Correlation with values of 3.61, 2.86 and −0.33 respectively, among all the models, as seen in **Table 1**. Our model also has a correlation of 0.87 which is the same as that obtained by the regular 3D CNN model and the information-theoretic 3D CNN model. Scatterplots between true age and predicted ages for all models are shown in figures 4, 5, 6 and 7. Notably, our method reduces the number of required parameters by 1.88 times in the case of the 3D convolutional neural networks and by 14.73 times in the case of the 3D Bagnet model. We also observe that training the recurrent slice-based model on sagittal slices yields better results than coronal and axial slices as seen in Table 2.

**Figure 4.**
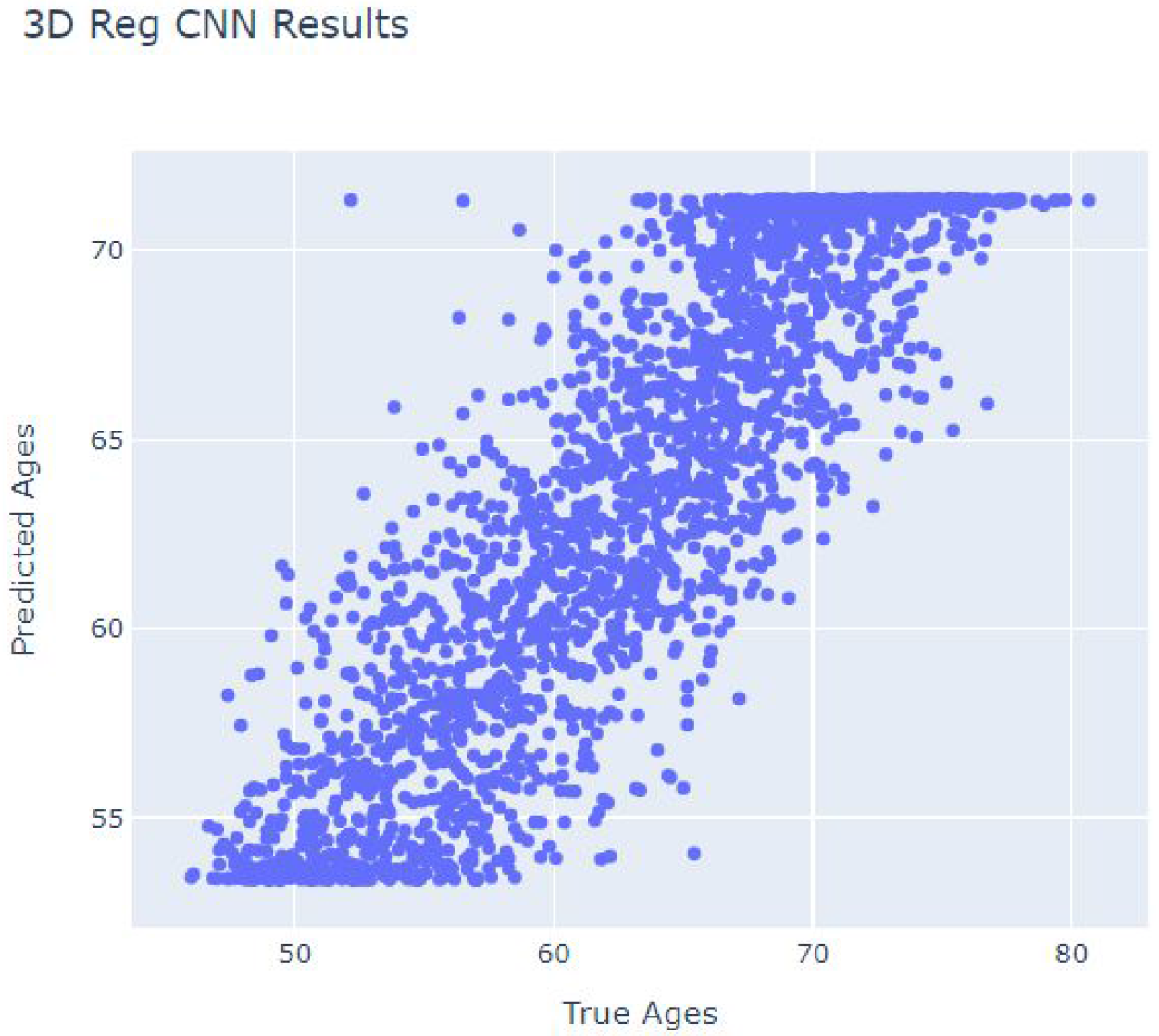
The scatterplot between True Age and the Predicted Age is shown for the 3D CNN Regression model on unseen data not used to train or tune the model.

**Figure 5.**
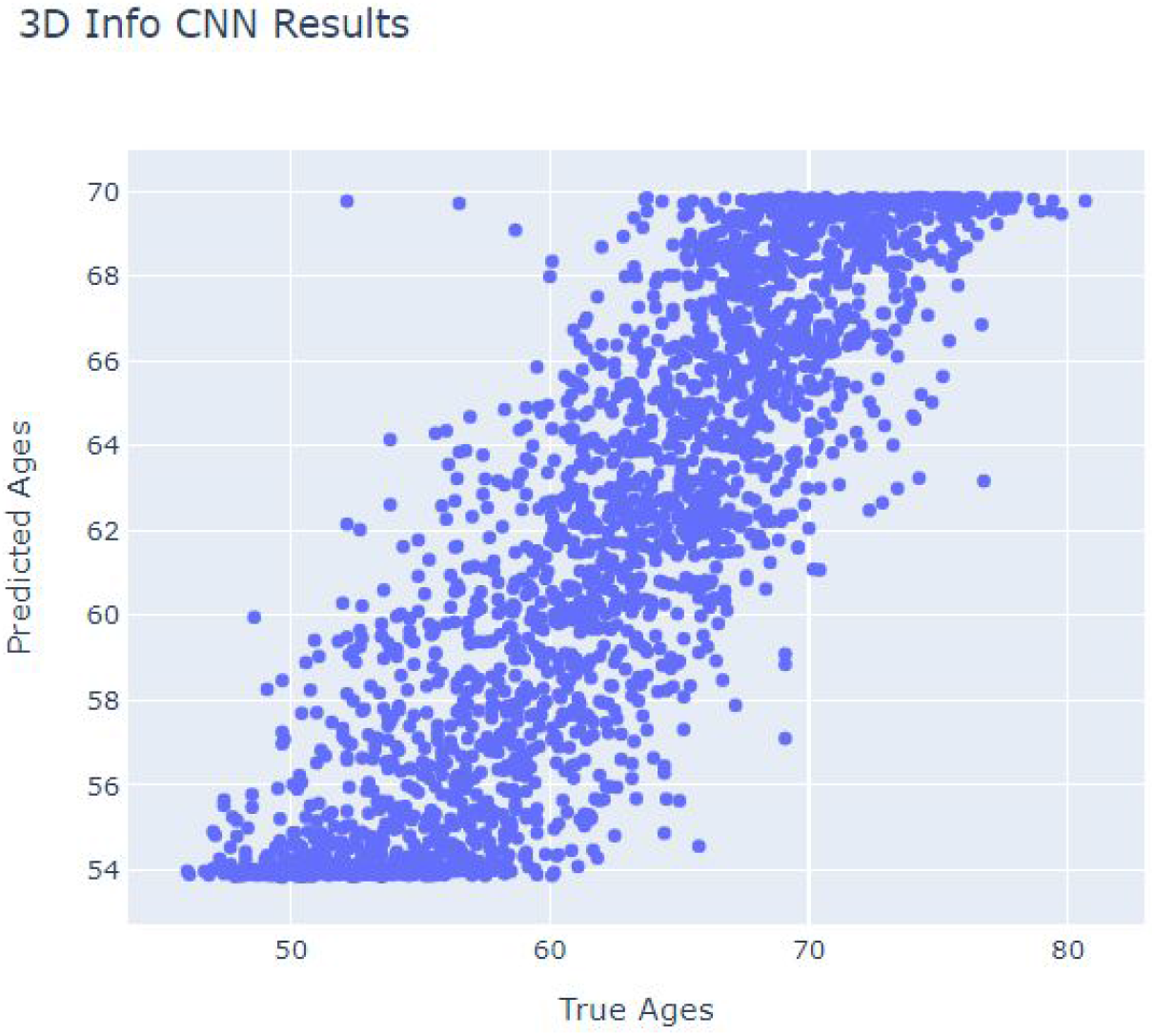
The scatterplot between True Age and the Predicted Age is shown for the 3D Info CNN model on unseen data.

**Figure 6.**
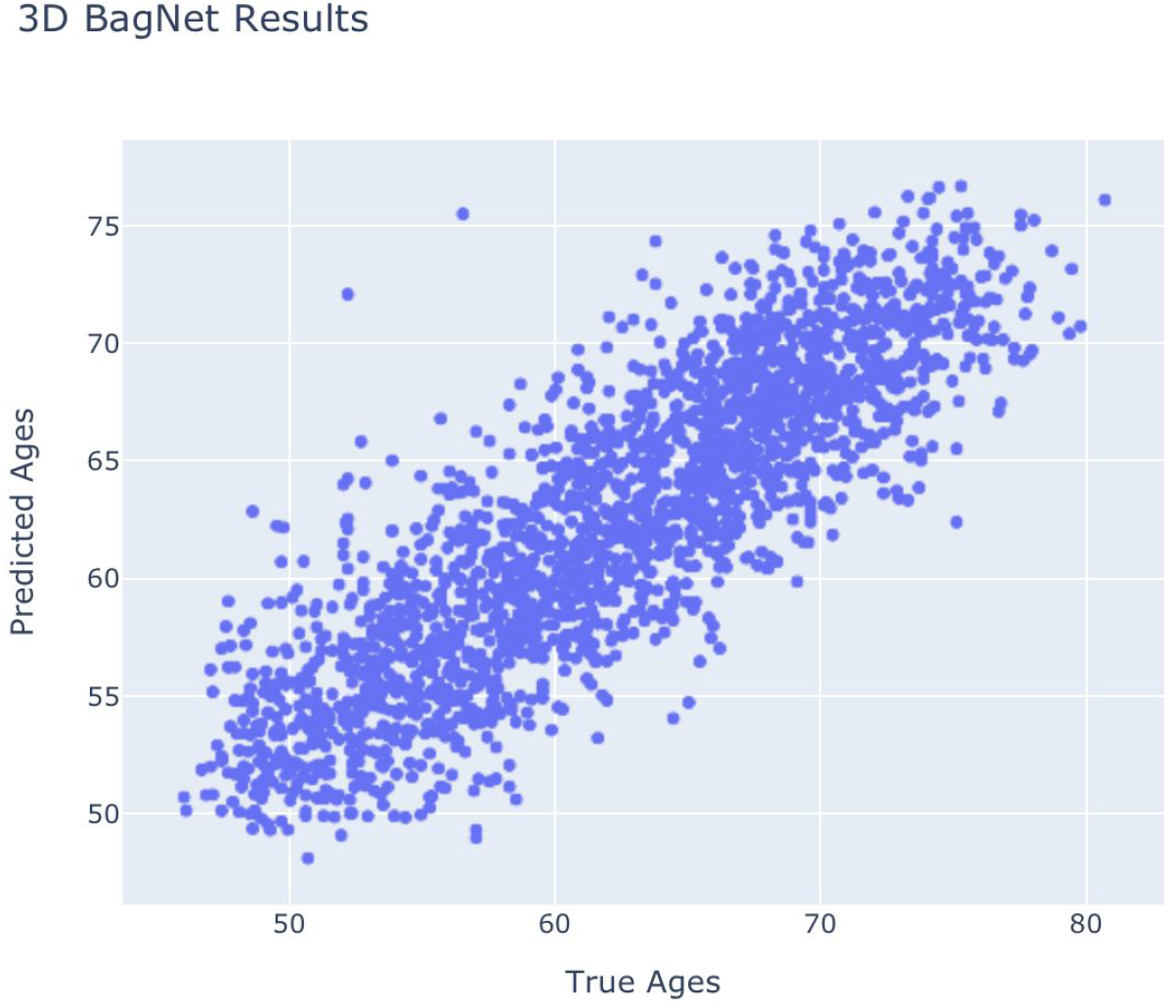
The scatterplot between True Age and the Predicted Age is shown for the 3D Bagnet model on unseen data.

**Figure 7.**
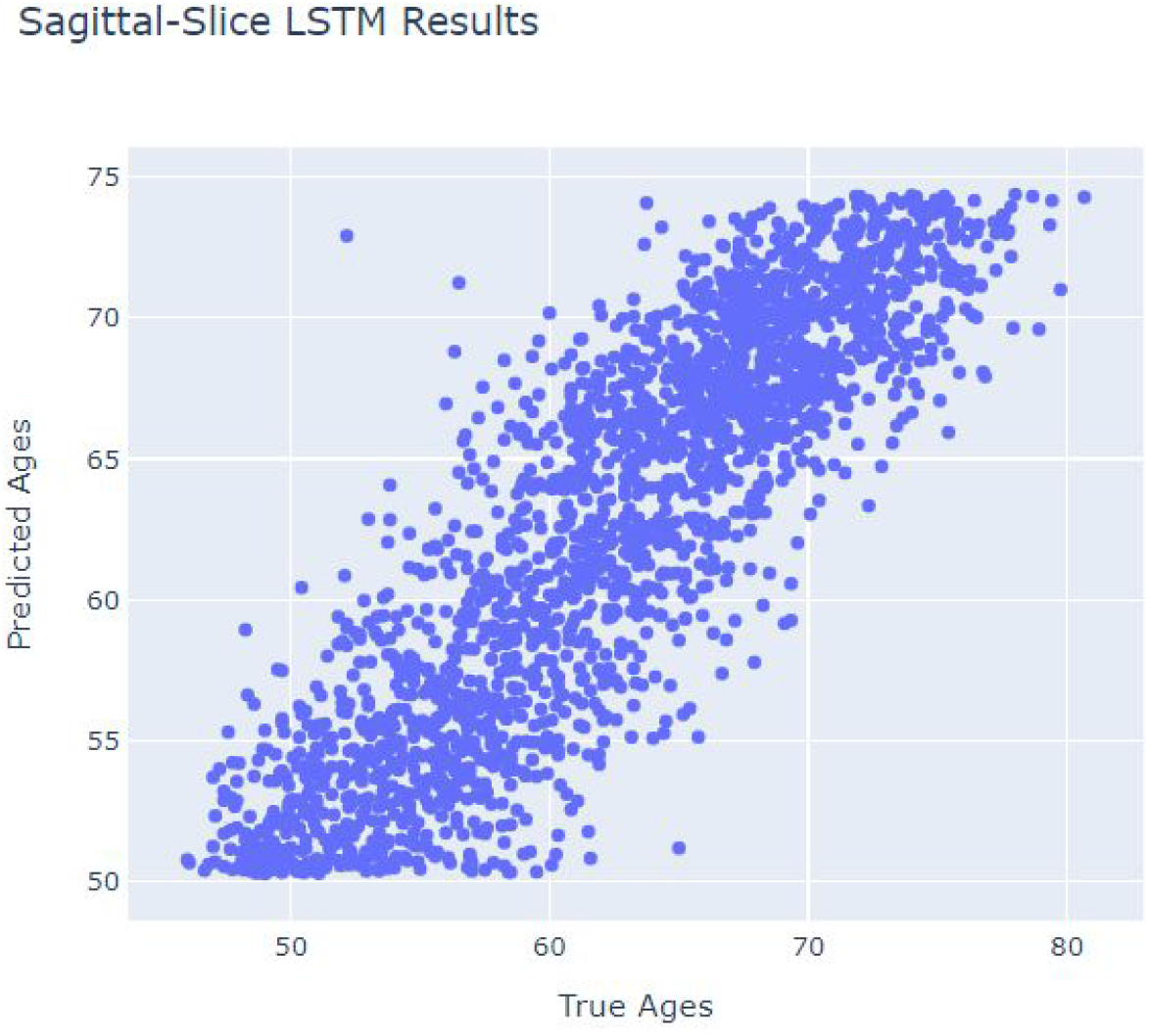
The scatterplot between True Age and the Predicted Age is shown for our 2D Sagittal Slice Sequence LSTM model on unseen data.

**Figure 8.**
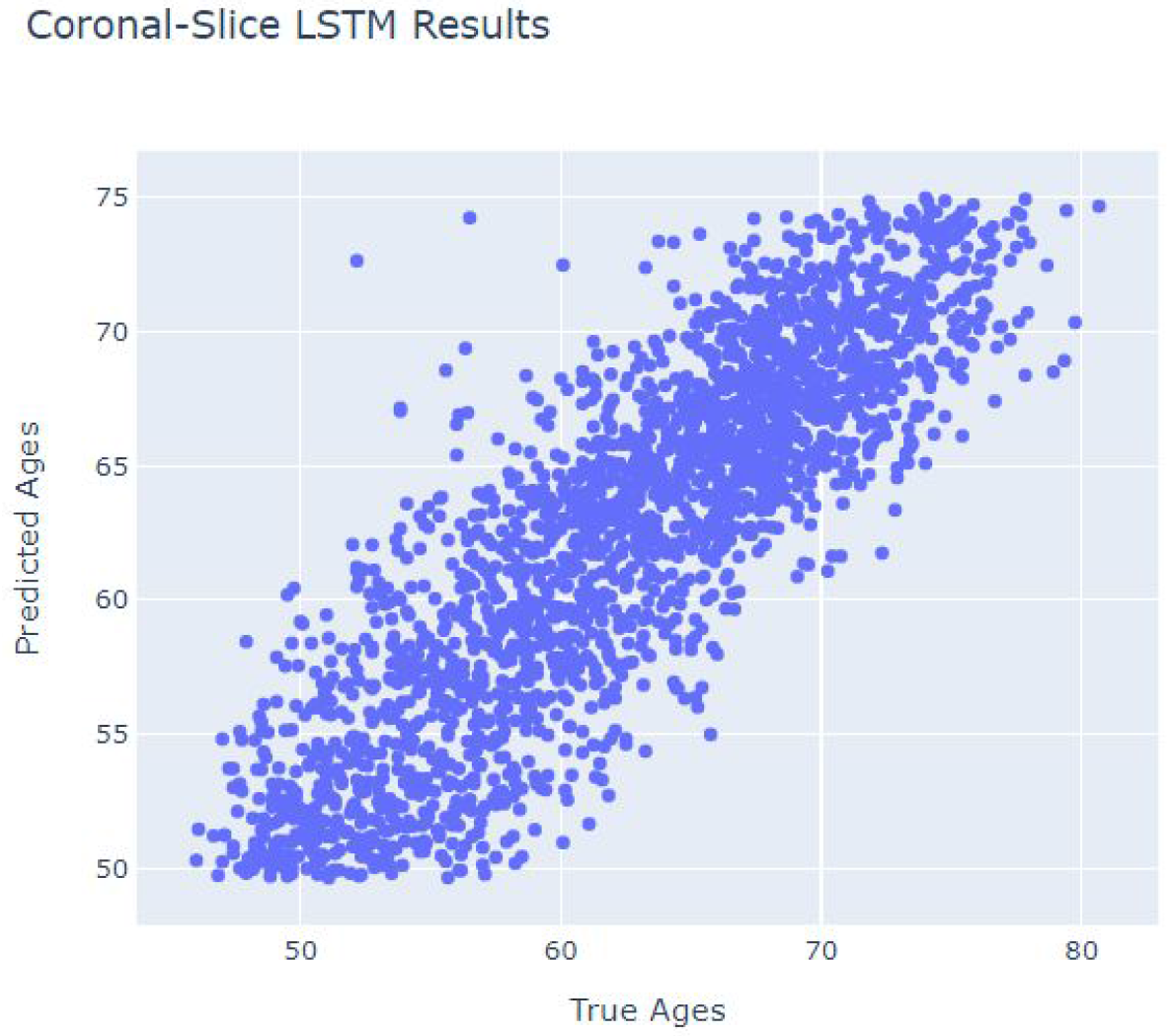
The scatterplot between True Age and the Predicted Age is shown for our 2D Coronal Slice Sequence LSTM model on unseen data.

**Table 2.** The results of our model on the three axes of MRI is presented here. When the model is trained on sagittal 2D slices, it performs better than when trained on data sliced along other axes.

|  | RMSE | MAE | Corr | Resid Corr |
| --- | --- | --- | --- | --- |
| Sagittal Slice | 3.61 | 2.86 | 0.87 | -0.33 |
| Coronal Slice | 3.76 | 2.98 | 0.86 | -0.44 |
| Axial Slice | 7.41 | 6.25 | 0.03 | -0.99 |

We note that there appears to be a strict upper bound for age predictions in Info, Regression, and slice-based LSTM models but not for BagNet. We stress that this is not by design and hypothesize that the large number of parameters (14 times higher than slice-based LSTM model) causes the BagNet model to learn more nuanced age distinguishing features. However, this may cause the BagNet model to overfit to training data.

## 4. DISCUSSION

While further and more rigorous comparison is necessary to compare model accuracy, our recurrent slice-based method performs comparably to other state-of-the-art methods, or better, while significantly reducing the required number of trainable parameters; it also achieves an MAE of 2.86 with only 1 million trainable parameters.

Our study has some limitations. In this initial report, we did not assess the correlations between BrainAge and neurological, psychiatric, and other health conditions. BrainAge - as with other biological clocks such as epigenetic age measures based on methylation of the genome - is considered by some as a measure of accelerated biological aging, so that the discrepancy between a subject’s true age and that estimated by the algorithm has been used as a predictor of mortality, health outcomes, and risk for disease. Even so, a recent report ^24^ identified a number of conceptual problems with this interpretation, noting that the brain age gap, as a residual, is dependent on age, so that any group differences on the brain age gap could simply be due to group differences in age. They further point to additional pitfalls in regressing out age from the brain age gap. Given these issues, we did not test associations here between our BrainAge method’s outputs and clinical outcome variables; in related work, however, Smith et al. ^11^ reported strong correlations of corrected brain age delta with 5,792 non-imaging variables (non-brain physical measures, life-factor measures, cognitive test scores, etc.), and also with 2,641 multimodal brain imaging-derived phenotypes, based on data from 19,000 participants in UK Biobank.

Future research will focus on a more rigorous comparison of our method with existing 3D deep learning approaches, studying the effect of transfer learning on the CNN encoder and on model training speed and accuracy. We will also adapt and evaluate the approach for other tasks such as disease classification, outcome prediction and prognostic modeling.

## ACKNOWLEDGMENTS

This work was supported in part by NIH grants R01 AG060610, R56 AG058854, U54 EB020403 from the Big Data to Knowledge (BD2K) program, R01 AG059874, P41 EB015922, and a research grant from Biogen, Inc. (Boston, USA).

